# Assessing biological network dynamics: Comparing numerical simulations with analytical decomposition of parameter space

**DOI:** 10.1101/2022.08.31.506131

**Authors:** Kishore Hari, William Duncan, Mohammed Adil Ibrahim, Mohit Kumar Jolly, Breschine Cummins, Tomas Gedeon

## Abstract

Mathematical modeling of the emergent dynamics of gene regulatory networks (GRN) faces a double challenge of (a) dependence of model dynamics on parameters, and (b) lack of reliable experimentally determined parameters. In this paper we compare two complementary approaches for describing GRN dynamics across unknown parameters: (1) parameter sampling and resulting ensemble statistics used by RACIPE (RAndom CIrcuit PErturbation), and (2) use of rigorous analysis of combinatorial approximation of the ODE models by DSGRN (Dynamic Signatures Generated by Regulatory Networks). We find a very good agreement between RACIPE simulation and DSGRN predictions for four different 2- and 3-node networks typically observed in cellular decision making. This observation is remarkable since the DSGRN approach assumes that the Hill coefficients of the models are very high while RACIPE assumes the values in the range 1-6. Thus DSGRN parameter domains, explicitly defined by inequalities between systems parameters, are highly predictive of ODE model dynamics within a biologically reasonable range of parameters.

## 1 Introduction

As the sophistication and scope of experimental methods in molecular and cell biology continues to grow, there is an increased need to use this data to synthesize a coherent understanding at the systems level. Experimental methods are successful if they isolate and study a particular feature of a complex system in isolation. The need to understand function of complex systems “as a whole” by combining partial insights is a goal of systems biology [26]. One of the principal demonstrations of the fact that such a unified insight has been achieved is to construct a mathematical model.

In this paper we discuss dynamical models that are built to represent the behavior of networks. Networks are often constructed from experimental data by postulating pairwise interactions of chemical species, e.g genes, their products, proteins and other molecules. The methodologies for discovery of such connections have varied levels of reliability, but none of them can measure the interaction between multiple effectors, or the range of behaviors of the network under varying conditions. Mathematical models are asked to integrate the local pairwise interactions and make predictions about the network behavior in conditions that are not directly experimentally accessible [33].

The principal challenge for model construction and validation is parameterization. The network structure constrains potential dynamics but does not uniquely determine it. In fact, the behavior of the network in different conditions, perhaps embedded in different individual cells, may be different precisely because the underlying kinetic parameters have changed. While there are several methods to determine the structure of the network, it is very difficult to measure parameters, especially because they may depend on precise experimental conditions. Because of this, even comparing a model prediction to an experiment is challenging; if the parameters for the experiment do not agree with the parameters of the model when simulated, a correct model may give a disappointing fit. On the other hand, having a good fit does not guarantee that the model is generalizable beyond the conditions that have been fit. Therefore, validation of a model should require that the description of the behavior of the model is provided not only for particular set of parameters, or for a particular initial condition, but includes a broad range of potential dynamics across parameters and initial conditions.

In this paper we compare two different approaches to describing a range of potential dynamics of a network. RACIPE [21] relies on random but judicious sampling of parameters and initial conditions with an ODE (Ordinary Differential Equation) based simulation that describes the interactions in the form of Hill functions, while DSGRN [5, 6, 15, 27] uses combinatorial computations to analyze all multi-level Boolean models compatible with the network dynamics. DSGRN embeds [4] discrete Boolean models into a continuous framework of switching systems [31, 32, 17, 18, 7, 11, 29, 22], which then permits the use of ideas from bifurcation theory to understand changes in dynamics as a function of parameters. This close relationship between Boolean and ODE descriptions leads to rigorous mathematical results that link dynamics described by DSGRN and that of smooth ODE systems with sufficiently steep nonlinearities [14, 10]. There are also explicit results for the size of allowed perturbations of equilibria predicted by DSGRN to systems with ramp nonlinearities [9]. DSGRN has been used successfully to describe complex dynamics of networks ranging from cell cycle [16] to EMT network [34]. RACIPE has been used to describe the dynamics of networks of varying sizes and biological contexts to understand the role of network topology in leading to various emergent phenotypes [19, 20, 24]. However, the role of parameters in the emergence of various phenotypes remains to be understood.

In this paper we compare RACIPE and DSGRN approaches to study dynamics of gene regulatory networks (GRNs). In Section 2.1 we use RACIPE to generate parameter samples and simulate dynamics of three two-node networks: Toggle Switch (TS), Double Activation (DA) and Negative Feedback loop (NF). We attempt to find parameters of the network that would predict behaviors like monostability vs. bistability. We do not observe any clear associations between individual parameters and GRN dynamic behavior. We hypothesise that the reason is that such behaviors depend on combinations of parameters, rather than on a single parameter. Since a combinatorial explosion prevents us from testing predictions based on all possible combinations of several parameters, we turn to DSGRN methodology. There is a direct translation between RACIPE parameters and DSGRN parameters, with the exception of the Hill coefficient *n* that is a parameter in RACIPE, but is missing in the DSGRN switching ODE model, since this model corresponds to the RACIPE model in the limit *n* → ∞. However, DSGRN provides explicit decomposition of the parameter space (**Fig 4**) into domains that have invariant dynamical behavior, which is directly computable without use of ODE simulations.

We are therefore able to compare RACIPE simulations to DSGRN predictions by locating the RACIPE parameter within a particular DSGRN parameter domain. We find a very tight fit for when the RACIPE samples Hill coefficients from range 10 − 100. Perhaps surprisingly, we also find that this fit does not deteriorate much when we sample Hill coefficients from range 1 − 10. Since this lower range is biologically more plausible, this suggests that the DSGRN parameter domain decomposition predicts dynamics for the biological range of Hill coefficients *n*.

We then test the close match between RACIPE simulations and DSGRN predictions on a three-node network Toggle Triad (TT). We find the same above mentioned good agreement.

Finally, we test an ensemble agreement between these two approaches. To do this we ask whether the frequency of dynamical behavior pooled across all parameter samples of RACIPE matches the frequency of dynamical behavior pooled across all parameter nodes of DSGRN. We find disagreement that persists for all values of *n*. We are able to fully explain this disagreement as caused by a non-uniform sampling of individual DSGRN parameter domains by RACIPE. In particular, for the examined two-node networks, the DSGRN parameter node that predicts bistability is sampled much more often than the nodes that predict monostability. When corrected for the nonuniform sampling, the agreement between DSGRN predictions and RACIPE simulations is restored.

By integrating the ideas from RACIPE and DSGRN, our results provide a way to deepen our understanding of the boundaries in the parameter space between distinct dynamical behaviors.

## 2 Results

### 2.1 Analysing RACIPE data to identify parameter phase boundaries

Using RACIPE, a parameter-agnostic approach that estimates the steady states of gene regulatory networks over a large parameter space, we aim to understand the regions of parameters that lead to a particular dynamical behavior of the network. Each parameter set specifies a system of differential equations whose dynamics in the phase space exhibits a long term behavior like steady states, periodic oscillations and multistability. In order for these behaviors to be observable, they must be stable, i.e., they must attract nearby initial conditions. We will use the word “phase” to denote different types of stable behavior of a system. Therefore the goal is to describe for each network a collection of attainable phases together with description of parameter domains that parameterize systems with that phase (See Methods).

We start by analysing a simple gene regulatory network called the Toggle Switch (TS): a network with two nodes and two edges, or equivalently “links”, such that each node inhibits the production of the other (**Fig 1a**). In the context of GRNs, nodes can be transcription factors or RNA molecules. We simulated TS using RACIPE, obtaining a map between parameter sets and the corresponding steady states (**Fig 1b**). We discretized these steady state values for ease of analysis, so that each steady state belongs to one of the four categories: high-high (11), high-low (10), low-high (01) and low-low (00). High and low levels of RACIPE steady states are defined based on whether the steady state level is higher (1) or lower (0) than the ensemble mean, where the ensemble is identified by the collection of parameter sets sampled by RACIPE. Simulating TS using RACIPE gives us two types of behavioral information: a) the number of steady states emergent from each parameter set (**Fig 1c**), and b) the category of the steady state(s) given by each parameter set (**Fig 1d**). The dominant dynamical behavior is monostability, with one node at a high level of expression and the other node low [21].

**Figure 1:**
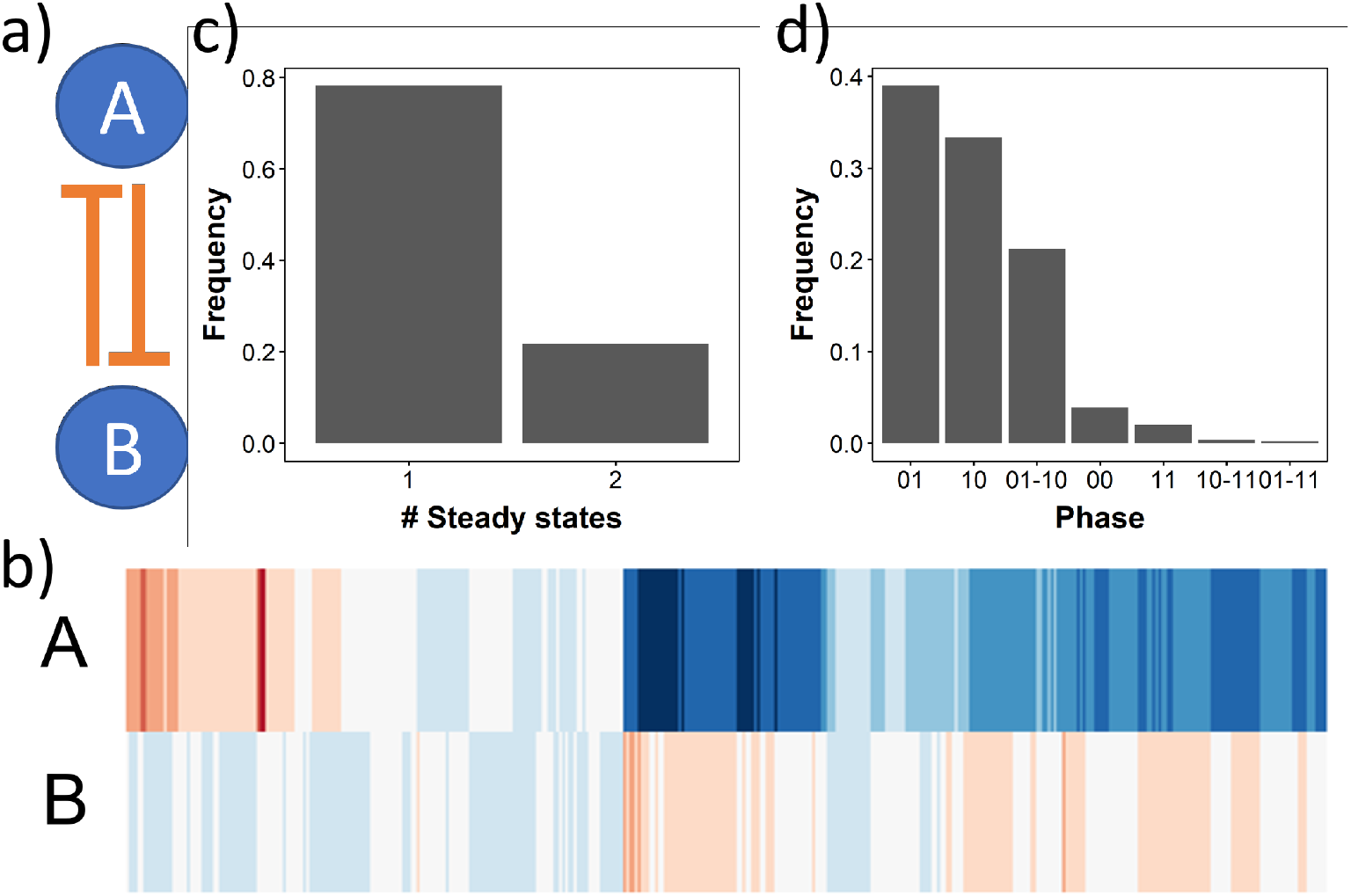
Output of RACIPE for Toggle Switch (TS) network. a) TS network structure. b) Heatmap representing all solutions from all parameter sets obtained via RACIPE. Red color indicates high, white indicates moderate and blue indicates lower expression levels of the node variable labelled to the left. c) Frequency of monostable (x-axis label 1) and bistable (x-axis label 2) parameter sets sampled by RACIPE. d) Frequency distribution of individual phases obtained from RACIPE for TS.

We then sought to understand the contribution of individual parameters in distinguishing between the categories of a given emergent behaviour. RACIPE samples five types of parameters (**Fig 2a**). For TS, which has two nodes and two edges, the parameters are labelled as follows

**Figure 2:**
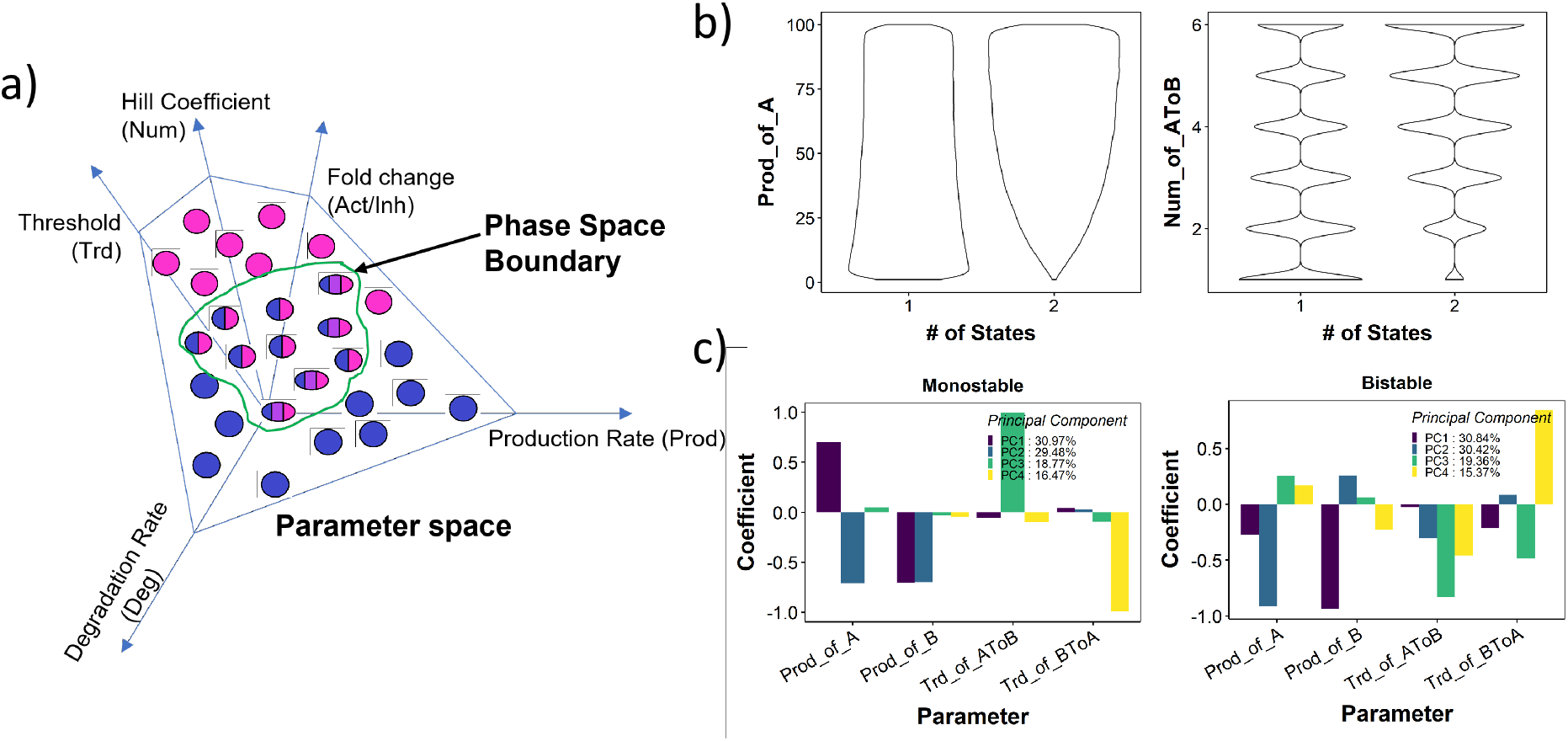
Identifying phase boundaries in RACIPE parameter space. a) Depiction of phase space boundaries in RACIPE. The parameter space is defined by the ranges of the five types of parameters. Each sampled parameter set can be monostable (unicolored ovals) or multistable (multicolored ovals). The phase boundary (green closed curve) separates monostable parameter sets from multistable parameter sets. b) Distribution of Production rate of A (left) and hill coefficient of A → B link (right) for monostable and bistable parameter sets in toggle switch. c) Barplot depicting the Principal component coefficients 1-4 for the four top parameters obtained from PCA of monostable (left) and bistable (right) parameter sets for toggle switch.

- Node level parameters
  - Prod of A(or B): Production Rate of node A (or B)
  - Deg of A(or B): Degradation Rate of node A (or B)

- Edge level parameters
  - Inh(or Act) of AToB(or BToA): Fold change
  - Num of AToB(or BToA): Hill coefficient
  - Trd of AToB(or BToA): Threshold for the link

The details of the ODE system are given in Methods section. We first checked if the distributions of any of the individual parameters can delineate monostability from multistability. We find that while there are no clear delineators, the bistable parameters show a lower frequency of low Hill coefficient, low production rate, and high degradation rate (**Fig 2b**, **Fig S1**).

We then performed PCA separately on monostable and bistable parameter sets. In both cases, the first four axes of PCA together could explain *>* 95% variance (**Fig 2c**, **legend**). At first glance, the primary contributors to the variance (highest coefficient in PCA axes) for both monostable and bistable parameter sets are the two production rates and two thresholds corresponding to the two edges in TS. Furthermore, in PC1 of the monostable parameter sets, the coefficients of the production rates have opposite signs, while the same coefficients in bistable parameter sets have the same sign. In PC2, the reverse pattern is observed, i.e., same sign in monostable parameter sets and opposite sign in bistable parameter sets. The threshold parameters had a stronger presence in PC3 and PC4 in both monostable and bistable parameter sets. However, while threshold parameters are exclusively limited to PC3 and PC4 in monostable parameter sets, they also contributed to PC1 and PC2 in bistable parameter sets. This indicates higher complexity of the description of the parameter set that exhibits bistability.

To better understand the patterns observed in PCA, we moved on to ask if the production rate and threshold value can delineate bistability from monostability. We generated a density plot of monostable and bistable parameters along the axes of both production rates, to identify the ranges of the production rates that are observed most frequently in mono and bistable cases. These plots indicated that while monostable parameter sets show higher incidence when two production rates are very different i.e. Prod of A *>>>* Prod of B and Prod of B *>>>* Prod of A, or when both production rates are low, in bistable parameter sets we see a higher incidence of approximately equal production rates Prod of A ≈ Prod of B (**Fig 3a**). Furthermore, the production rates in bistable parameter sets had a higher chance of being above the median production rate sampled by RACIPE (50). A similar visualization involving threshold parameters revealed that both threshold values tend to be lower than median and similar to each other in bistable parameter sets (**Fig 3b**). No clear trend was observed for threshold in monostable parameter sets, suggesting that threshold values being smaller is necessary but not sufficient condition for bistability.

**Figure 3:**
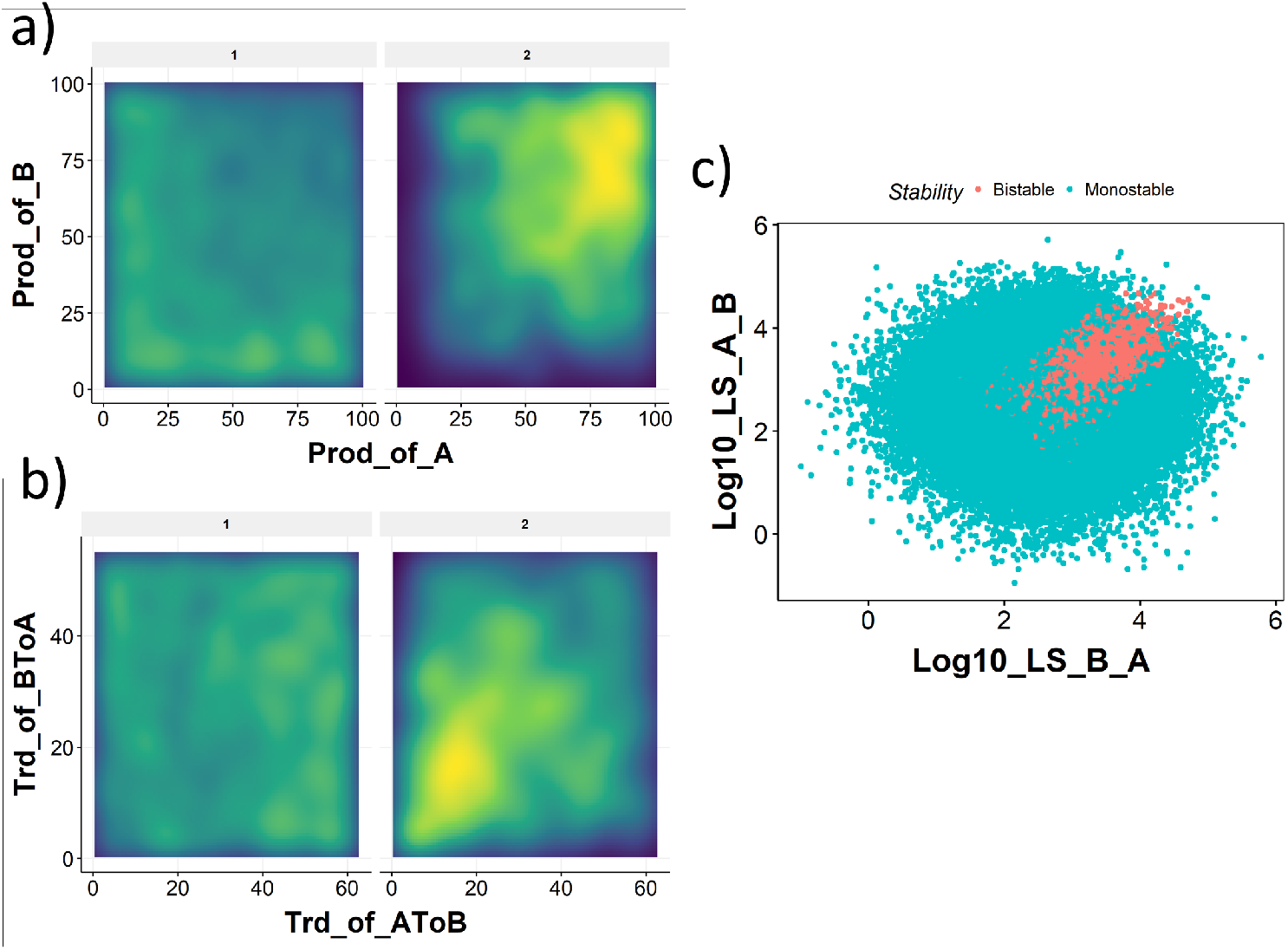
Link strength delineates monostable and bistable parameter sets. a) Density distribution of the parameters Production rate of A and B for monostable (panel label - 1) and bistable (panel label - 2). The color scale varies from purple (low) to green (medium) to yellow (high). Higher density indicates higher number of parameters in the corresponding region. b) Same as a), but for Thresholds for *A* → *B* and *B* → *A* links. c) Same as a) but for Log10 values of link strengths of *A* → *B* and *B* → *A* links. d) Scatterplot between the link strength values demonstrating the separation of parameter domains by link strength.

These patterns can be interpreted as follows: high production rate of A and low threshold from A to B implies a higher chance of the inhibition from A to B being active; similarly, a high production rate of B and low threshold from B to A implies a higher chance of the inhibition from B to A being active. Hence, we hypothesized that occurrence of bistability in TS requires both links to be simultaneously active. On the other hand, the emergence of monostability is associated with asymmetry of the production rates, where one of them is higher than the appropriate threshold, while the other one is lower than its threshold. Therefore we hypothesize that monostability occurs when one of the links is significantly stronger than the other, while bistability occurs when both links have similar and high strength.

To test this hypothesis, we adapted a measure of link strength [8] to explain the emergence of certain phenotypes in GRNs. The link strength takes the following non-dimensional form:

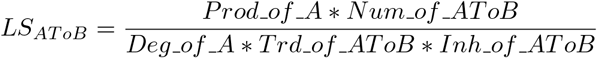

We generated a scatter plot of the log value of the link strengths on either axis (**Fig 3c**). Interestingly, using a logarithm transformation on these two parameters converts the set of sampled parameters into a circular cloud where parameters that support bistability form a small section of the cloud. Note that this section is in the region that corresponds to both links being strong and similar in value, which is consistent with our hypothesis that both links should have similar and high strength for bistability.

We further tried to delineate different phases (10, 01, 01-10 etc) of mono and bistability using link strength S2. Low link strength (*<* 3) for both links leads to 00 phase. For a majority of the parameter sets displaying phase 10, link strength of A to B is low. Similarly for phase 01, link strength of B to A is low. However, we find it hard to distinguish between 11 and 01-10 phases using link strength analysis. Furthermore, for phases 10 and 01, there exist cases with high link strength for both links.

While link strength can delineate the monostable and bistable parameter sets better than any individual/combination of parameters tried so far, the boundary between bistability and monostability and that between individual phases is not completely sharp. This leads to uncertainty in prediction of multistability of a parameter for TS. Therefore we tried to clarify and sharpen this boundary through DSGRN.

### 2.2 DSGRN inequalities define clear boundaries separating phases

DSGRN (Dynamic Signatures Generated by Regulatory Networks) is a modeling platform that assigns to a GRN a *switching ODE system* with undetermined parameters [5, 6, 15, 27]. The parameters include a threshold value, as well as low and high production rates assigned to each edge and a decay rate assigned to each node. At each node, if the combination of high and low values transmitted along the incoming edges is higher than a particular threshold of an outgoing edge, this edge is activated. In the case where this is an activating edge, a high production rate is triggered; when there is a repressing edge, a low production rate is triggered. Using these parameters, DSGRN provides an explicit finite decomposition of the parameter space into parameter domains, such that for all parameters in each domain the state transition graph (STG) (see Methods section for detailed description), the phase is invariant.

The parameters of the switching ODEs that are used by DSGRN can be related to the parameters used by RACIPE formalism (Methods 4.5). We therefore asked if DSGRN inequalities can be used to delineate parameter spaces in RACIPE and predict the outcome of the dynamics at parameters sampled by RACIPE. As a first step, we carefully compared the structure of the ODE models used in RACIPE and DSGRN (see Methods).

There are two major differences between RACIPE and DSGRN. First, while RACIPE uses model equations with a finite Hill coefficient (sampled by default between 1 and 6), DSGRN uses piece-wise constant nonlinearities that can be obtained as a limit of Hill functions where the Hill coefficient *n* → ∞. Second, in DSGRN the production rate for a node is calculated independently for each incoming edge, followed by combining these production rates together (taking a product) to get the net production rate of the node. In RACIPE, each node is assumed to have a basal production rate which gets multiplied by a fold change parameter for each incoming edge. Assuming that each incoming edge has an equal contribution to the basal production rate, we establish a one-to-one correspondence between the parameters of RACIPE and DSGRN, with the exception of the Hill coefficient. With this translation between these two approaches, we find that the link strength formalism described previously has a similar form as the inequalities obtained from DSGRN that define the parameter nodes.

DSGRN does not predict what phase will be attained for a given initial condition or specific real-valued parameter set. However, an explicit switching ODE system can be constructed that does and faithfully reflects the predictions of DSGRN, see Methods 4.5. This framework allows a direct comparison between a RACIPE simulation and the corresponding DSGRN prediction for a set of initial conditions and parameters for a given GRN. “Switching system” and “DSGRN” may be used interchangeably in what follows.

Using this correspondence we imposed the inequalities calculated in DSGRN onto RACIPE parameter sets, and obtained a phase distribution for each parameter domain, where a parameter domain is defined by a unique combination of inequalities (**Fig 4**). Because DSGRN corresponds to a RACIPE model where the Hill coefficient *n* approaches infinity, i.e. for very steep Hill nonlinearities, we expect that as *n* gets larger, the correspondence between DSGRN prediction and RACIPE simluation will improve. However, this argument does not provide any information on the accuracy of this prediction for relatively small values of *n*. Somewhat surprisingly, most of the monostable parameter nodes show nearly identical behavior between RACIPE simulation and DSGRN prediction, even at relatively small values of Hill coefficient *n* ∈ {1, 舰, 10}. However, at low values of *n* the central parameter node (parameter node 4, **Fig 4**) shows larger discrepancies. For these sets of parameters DSGRN predicts bistability between steady states 01 − 10, i.e., any parameter set obeying the inequalities will exhibit bistability with the states being 10 and 01. RACIPE on the other hand shows a more heterogeneous phase distribution, with predominant phase still being bistable 01 − 10, but also registering monostable states 10 and 01 (**Fig 5** center panel, parameter node 4).

**Figure 4:**
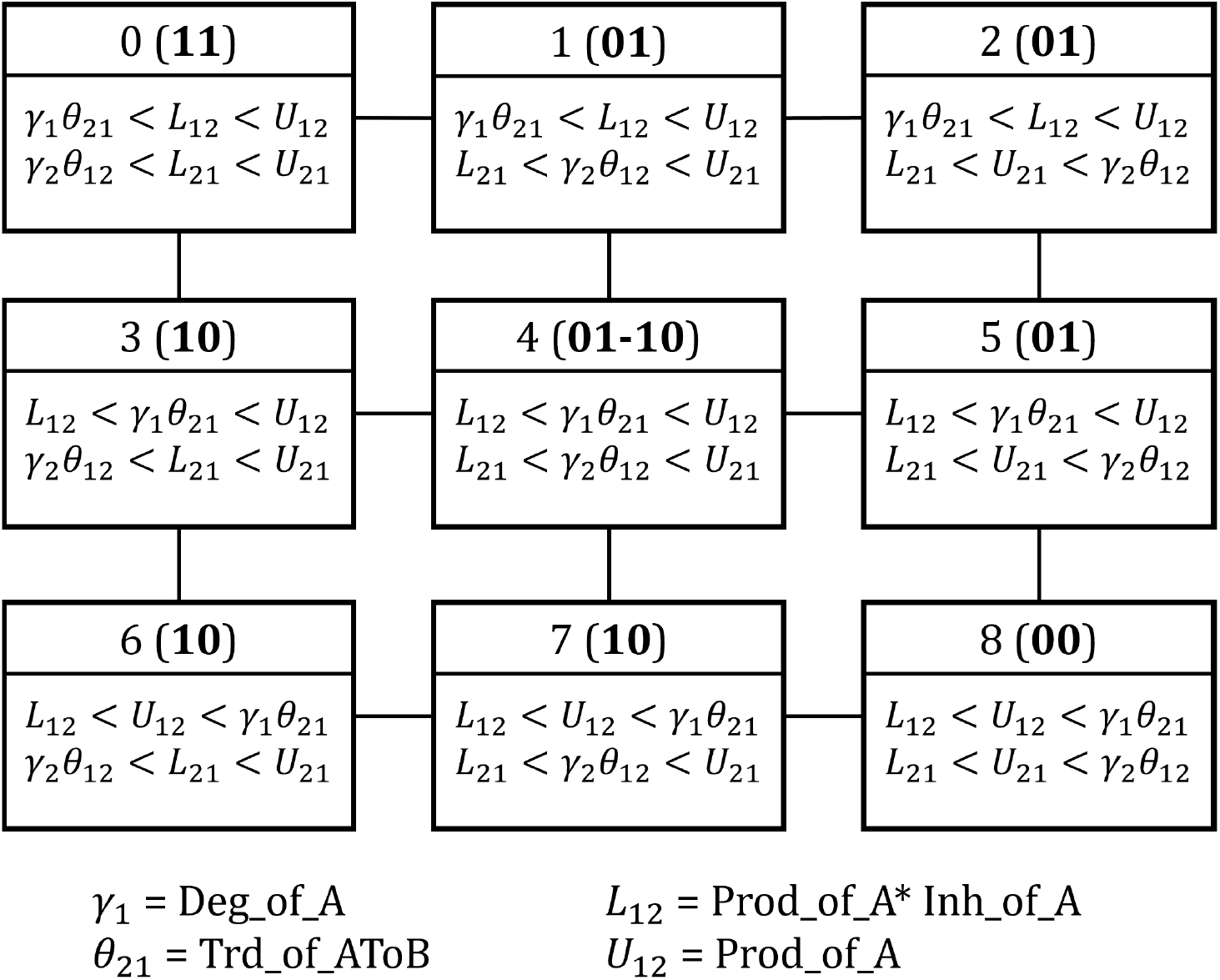
Depiction of the parameter domains/nodes calculated by the DSGRN inequalities for TS. Each box represents a parameter node. The inequalities that define the parameter node are given in the lower part of the corresponding box. At the top of each box, the integer provides a reference to each parameter node, followed by the description of the phase at each parameter sample satisfying the inequalities, as predicted by DSGRN.

**Figure 5:**
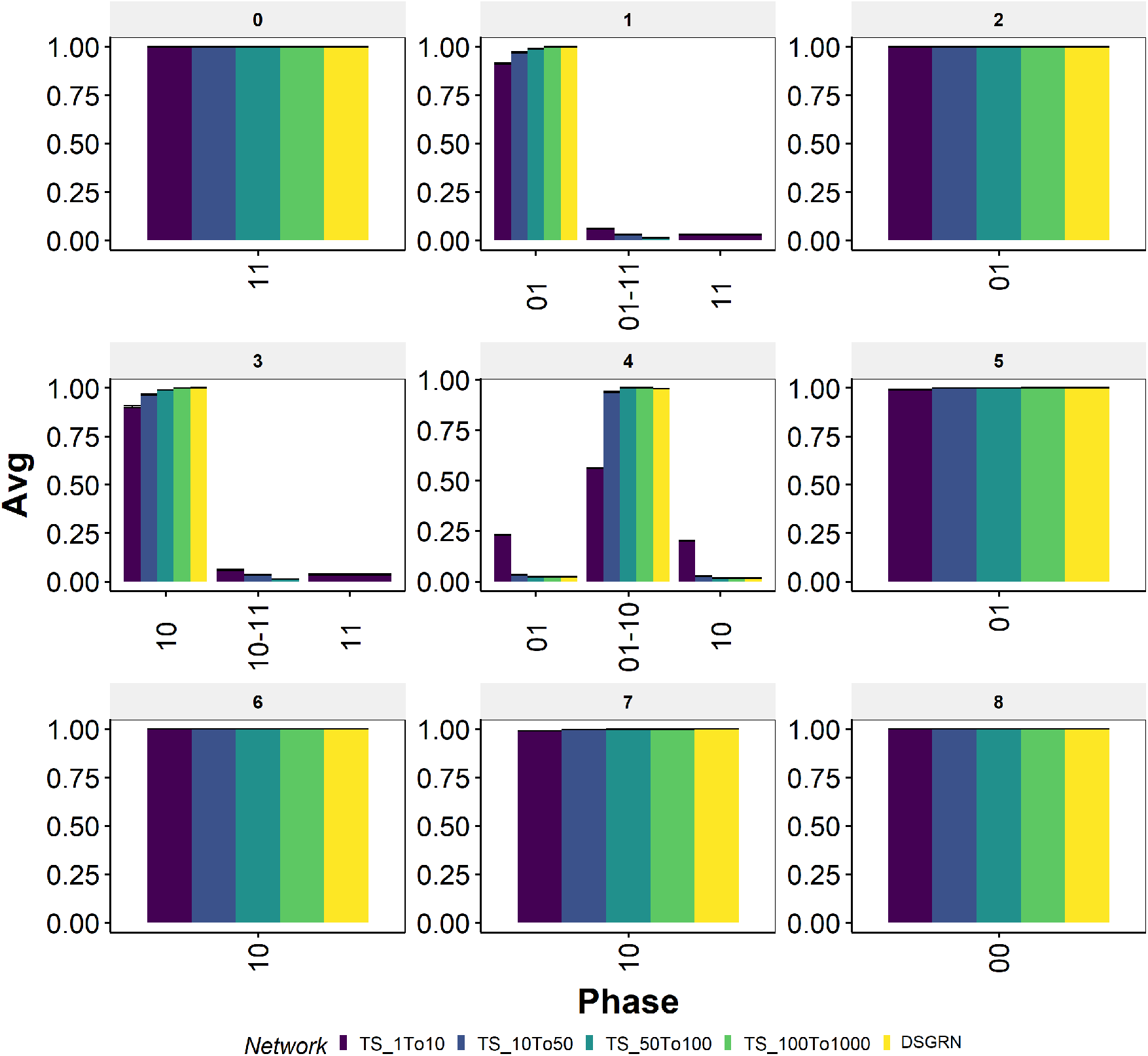
Phase distribution of switching system against different ranges of Hill coefficients in RACIPE for TS. The switching system is represented by yellow colored bars. Default RACIPE conditions are represented by the dark blue colored bars. The range of Hill coefficients in each case is reported in the color-legend.

To check whether the increase in the Hill coefficient will bring the RACIPE simulations and DSGRN predictions closer together, we modified the RACIPE parameters by sampling Hill coefficient values uniformly from various ranges (1 − 10, 10 − 50, 50 − 100, 100 − 1000) and compared the parameter node-wise phase frequencies with those of DSGRN (**Fig 5**). While the Hill coefficient range of 50 − 100 shows near identical results as that of DSGRN, even range of 10 − 50 is similar to DSGRN within reasonable error (*<* 5%). Further probing of parameter node 4 in RACIPE revealed that, having both Hill coefficients greater than 5 is enough to get the frequency of bistable phase (10-01) to be greater than 90% (**Fig S3**).

We observed similar trends for the other two node network motifs: Double Activation (DA) and Negative Feedback loop (NF), (see **Fig S4**, **S5**, **S6**). For both of these networks, RACIPE shows similar results as DSGRN at moderately high Hill coefficient values (*>* 10). Therefore, DSGRN inequalities can be used to clearly delineate RACIPE parameter space into different phases. Furthermore, DA network shows bistability for parameter node 4 (FP(00-11)), which for some parameter sets is lost in RACIPE at lower values of Hill coefficient, but s recovered with increase in Hill coefficient values.

NF shows particularly interesting trends. The “frustration” in NF network (i.e., while A activates B, B inhibits A, causing the state of the system to oscillate [3]) leads to the prediction of cycles in DSGRN at parameter node 4. For the corresponding parameters, RACIPE at lower values of Hill coefficients (*<* 10) predicts predominant monostability. We wanted to check if any of the parameters have been mislabelled as monostable while they are actually oscillatory, since RACIPE cannot identify oscillations. First, we confirmed whether the output of RACIPE is actually a steady state by imposing the condition that the derivative of all nodes should be small for steady states (see Methods section) We find that for all parameters predicted by DSGRN to be cyclic, the steady state obtained from RACIPE satisfied the derivative condition with a tolerance value of 10^*−*4^. Interestingly, as we increased the range of Hill coefficient to 10 − 50, RACIPE predicts all of these parameter sets to have ten steady states, which is the maximum number of steady states RACIPE can detect for a given parameter set. None of these states have a low derivative. Because DSGRN predicts cycles for these parameter sets, we categorised the parameter sets that show ten states in RACIPE, with none of them being a steady state, as cyclic.

The analysis so far suggests a transition from monostability to multistability (cyclic behavior for NF) in some RACIPE parameters as the Hill coefficient increases, a frequently observed pattern in GRNs [2]. This observation implies that the parameter sets that showed monostability at low Hill coefficients can acquire a new behavior as the Hill coefficient increases. These new behaviors will be more or less detectable numerically depending on the relative size of their basins of attraction. It is desirable to have a prediction not only of phase type and number, but also the relative sizes of their basins of attraction.

An analytical computation of the volume of the basins of the attraction in bounded region of phase space even for a simple system like TS is intractable due to nonlinearities. However, the boundaries of the basins of attraction for the TS switching system can be analytically computed (Methods section 4.4). Then any initial condition can be analytically associated to its long-term phase by its relation to the basin boundaries. The fraction of sampled initial conditions that go to a given phase is then an estimate of the relative basin size for that phase. There is no similar analytic boundary calculation for RACIPE, but the relative basin size can be numerically approximated by taking the fraction of initial conditions that converge to a given phase. We use these approximations to compare RACIPE basin sizes with various Hill coefficients to the limiting behavior in the switching system.

For comparison purposes, we defined a scalar “basin strength” to be the product of the fraction of initial conditions that converge to each detected phase. For monostability, this fraction can be at most 1 and for bistability it is maximally 0.25 (0.5 * 0.5 for equal basin sizes). We then calculated the difference between the RACIPE basin strength and DSGRN basin strength. The difference can range between −1 and 1, such that −1 is attained for parameter sets where RACIPE predicts monostability (basin strength 1) while DSGRN predicts an uneven bistability (i.e., low value of basin strength). Since the analysis is carried out for parameter sets belonging to parameter node 4, the basin strength for DSGRN will always be less than 0.25, making the upper limit for the difference 0.25.

In **Fig 6**, we plot this difference for a collection of 10000 randomly chosen parameters for both high and low Hill coefficients. We observe that for a given parameter set, as the RACIPE Hill coefficients increase, the basin strength of RACIPE simulations get closer to that of DSGRN basin strength (left and middle panels). This means that DSGRN boundary computations can be used to predict the relative sizes of basins of attraction.

**Figure 6:**
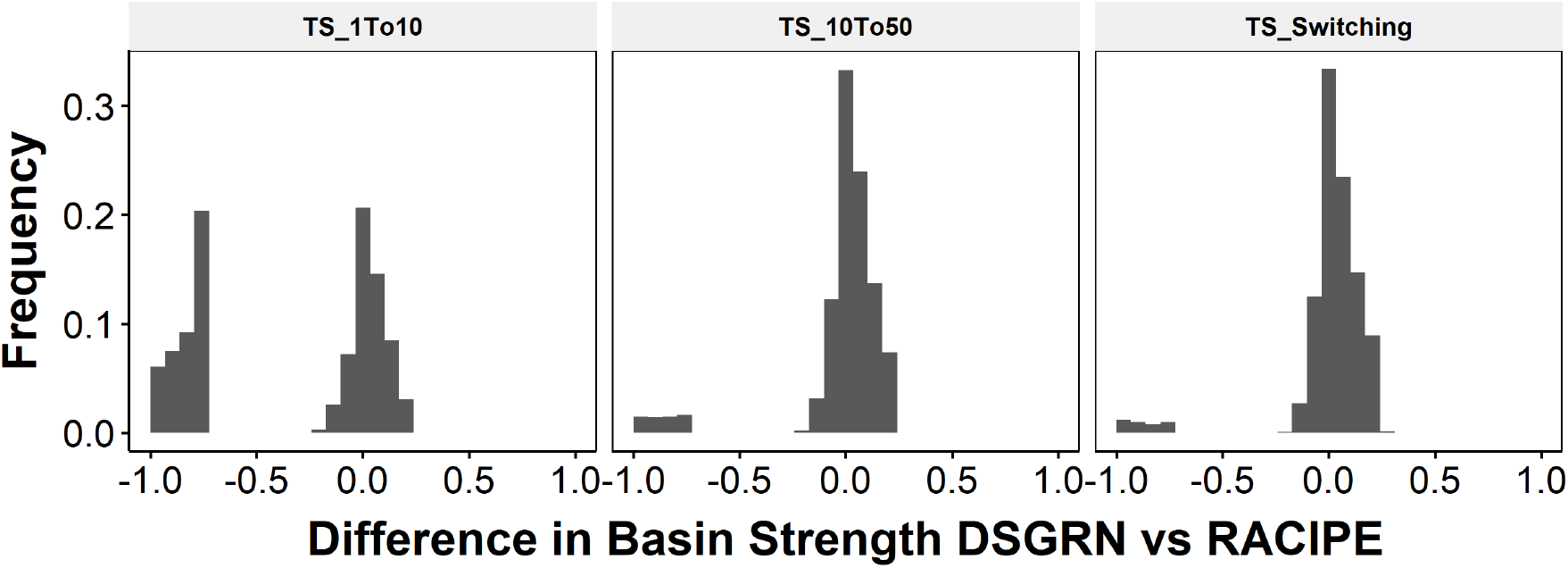
Switching system and RACIPE at high hill coefficient show similar basin strengths as DSGRN.

In larger networks for which analytical boundary computations in DSGRN are infeasible, an estimation of basin of attraction size for the switching system can be made analogously to RACIPE by numerically computing the fraction of initial conditions that converge to each phase. In **Fig 6** (right), we compare the analytical assignment of phase via basin boundary to this numerical approximation. We see that such a switching system approximation identifies the correct phase in about 95% of cases, making the contribution of the numerical simulations to the error in prediction about 5%.

### 2.3 Similarity between RACIPE and DSGRN holds for Toggle Triad

So far, we have found that in all three two-node networks that we studied, there is close correspondence between RACIPE and DSGRN. This suggests that DSGRN inequalities are able to describe how the dynamical behavior of the RACIPE model depends on parameters and delineate parameter regions of different phases. We now examine if these results hold for larger networks. To do this we chose Toggle Triad (TT), a three node network that can be viewed as a coupling of three toggle switches with two embedded negative feedback loops (see **Fig 7**). The most interesting feature of TT is that it can exhibit tristability (High-Low-Low, Low-High-Low, Low-Low-High).

**Figure 7:**
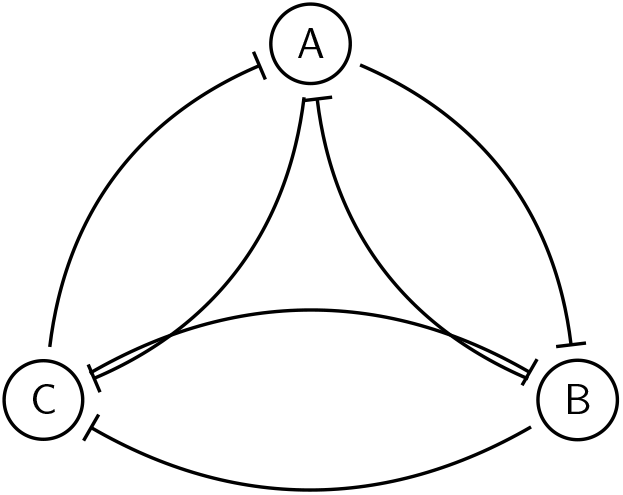
Toggle Triad (TT) network structure.

Since each node in a toggle triad has two inputs, the number of DSGRN parameter nodes is much higher than that for the two node networks. Therefore, instead of focusing on a particular parameter node, we decided to study the distribution of phases across RACIPE parameters sample sets. In **Fig 8a**, we compare phase predictions at each parameter given by RACIPE simulations to the predictions given by switching system simulations (equivalent to predictions by DSGRN). As with the two node networks, the similarity between RACIPE and DSGRN predictions increases with increasing Hill coefficient. Furthermore, we see a higher frequency of tristable phase 002-020-200 in DSGRN and in RACIPE with high Hill coefficients compared to RACIPE with lower Hill coefficients. A closer look at the parameter sets that are tristable in DSGRN reveals that greater than 30% of these parameter sets show tristability at low Hill coefficients in RACIPE. Of the remaining 70%, more than half show bistability equally distributed between 002-020, 002-200 and 020-200, which are all different subsets of the tristable phase, see **Fig 8b**. To eliminate the possibility that at low Hill coefficients, the tristability is not detected due to a smaller number of initial conditions, we simulated these networks for an increased number of initial conditions. We found that for 1000 and 10000 initial conditions, we do not observe the missing steady states for any initial condition. The difference in basin strengths also vanishes between the switching system and RACIPE (**Fig 8c**), suggesting that these parameter sets that showed bistability at low Hill coefficient gain another steady state as *n* increases.

**Figure 8:**
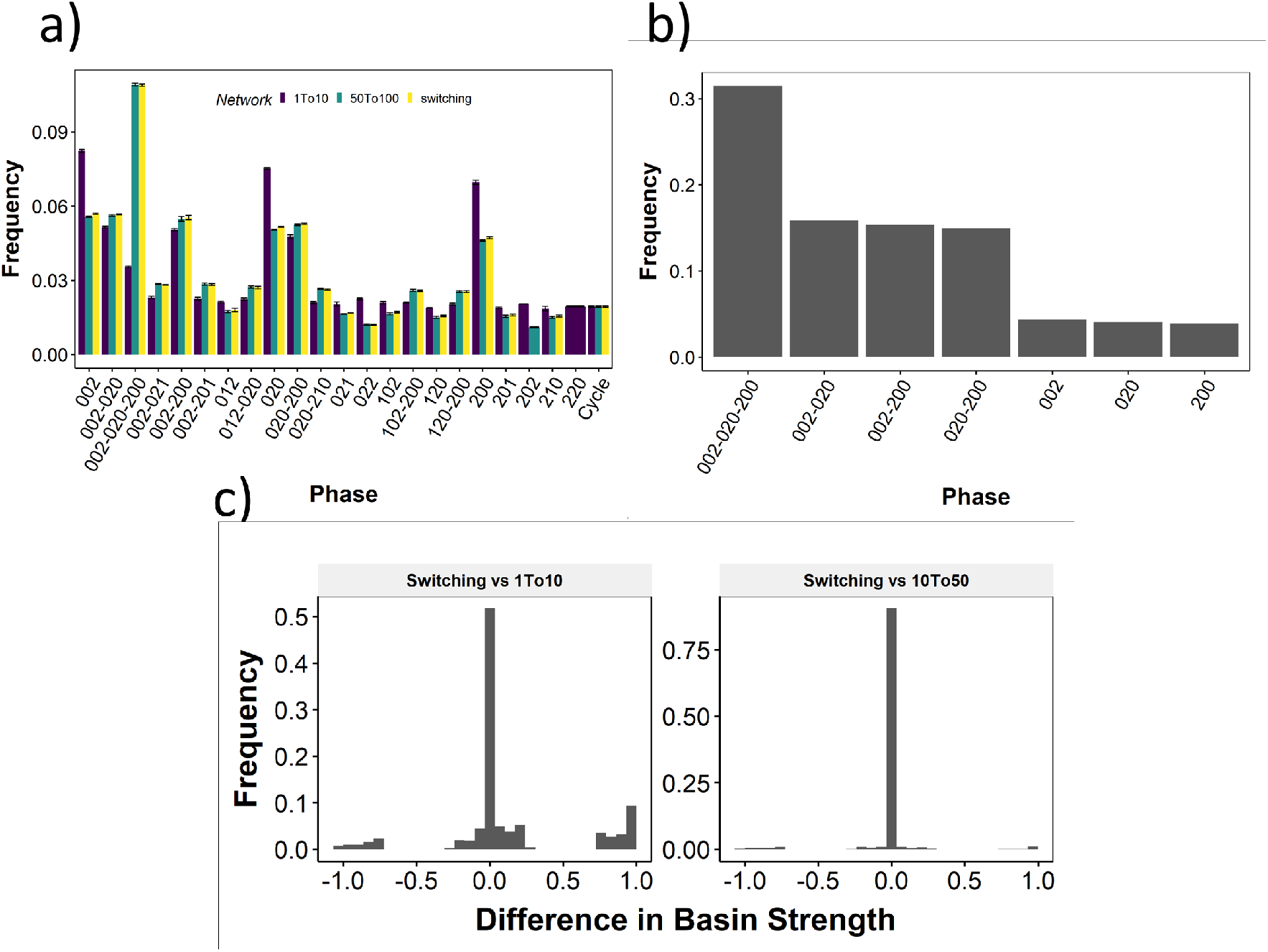
Comparison of RACIPE and DSGRN for Toggle Triad. a) Bar graph depicting the change in mean frequency of phase occurrence for RACIPE with Hill coefficient between 1 and 10 (purple), 50 and 100 (green) and switching system (yellow). b) Phase frequency distribution for parameter sets that exhibit tristability in switching system in RACIPE model with *n* ∈ [0, 10]. c) The difference in basin strengths for switching system and RACIPE with low Hill coefficients (right) and RACIPE with high Hill coefficients (left).

### 2.4 Contribution of the parameter sampling in RACIPE to the differences in RACIPE and DSGRN

Both RACIPE and DSGRN are capable of predicting the ensemble level behavior of a GRN. Given the similarities between RACIPE and DSGRN formalisms at high Hill coefficient, we compared the ensemble distributions, i.e. frequency distribution of individual phases obtained across all sampled parameter sets for RACIPE and all parameter nodes for DSGRN for the 2-node networks TS, DA and NF. Unlike the comparisons across individual parameter nodes, the ensemble frequency distribution of RACIPE predicted a higher frequency for the 01 − 10 phase as compared to DSGRN (**Fig 9**a). As the differences in the model formalism should diminish at high Hill coefficient values, we looked at the distribution of parameters sampled by RACIPE with respect to the DSGRN parameter domains. Interestingly, we find that RACIPE’s sampling method is highly biased towards the parameter node 4, which explains the prediction of high bistability from RACIPE at high Hill coefficient **Fig 9b**). Importantly, while DSGRN parameter domains decompose the parameter space into disjoint number of parameter domains (for TS, DA and NF there are 9 domains), this method is agnostic on where the biologically relevant parameters lie. If we assume that the RACIPE methodology samples biologically relevant parameters, then the non-uniform distribution of samples may be taken as a hypothesis which of the parameter domains are more important. The results here suggest, that even though only 1 out of 9 DSGRN parameter domains supports bistability, this domain is sampled by more than 50% of RACIPE parameters and hence it is more important than the ratio 1*/*9 would suggest. After normalizing the phase frequencies by the number of RACIPE parameters in each parameter node, the ensemble frequency distribution does match DSGRN predictions (**Fig 9c**).

**Figure 9:**
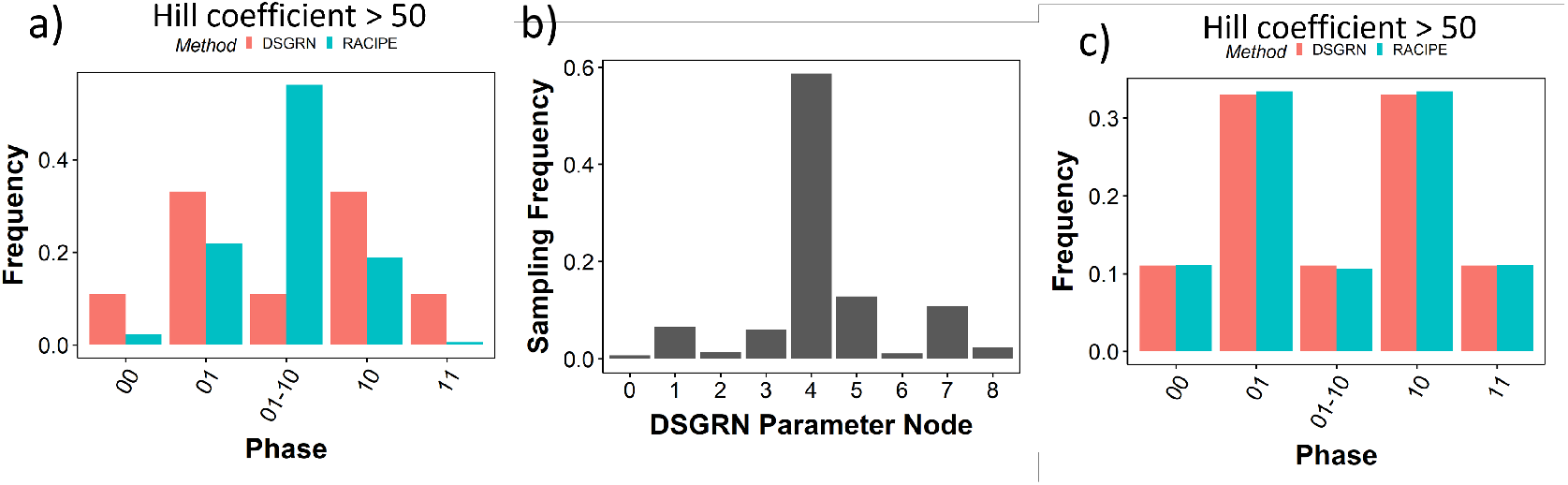
RACIPE shows uneven sampling of DSGRN parameter nodes. a) Comparision of phase frequency distributions for RACIPE with high Hill coefficient (green) and DSGRN (red). b) Distribution of DSGRN parameter nodes sampled by RACIPE. c) Same as a), but RACIPE phase frequencies are normalized by number of samples from corresponing DSGRN parameter domains.

With the default range of Hill coefficients 1 − 6, RACIPE’s ensemble distribution is much closer to that of DSGRN (**Fig S6**). We have seen before that RACIPE samples the bistable/cyclic parameter sets (parameter node 4) with higher frequency relative to the other parameter nodes of DSGRN. However, at low Hill coefficients, this bistability/cyclicity deteriorates to monostability for a significantly fraction of parameter sets belonging to parameter node 4. This deterioration serves to compensate for the biased sampling RACIPE conducts, thereby making the ensemble phase distribution of RACIPE at Low Hill coefficient similar to the corresponding DSGRN distribution.

## 3 Discussion

Modeling gene regulatory networks is qualitatively different than modeling physical systems like those in celestial mechanics and ecology [25]. While the behavior of the latter is described by equations stemming from physical laws and whose parameters are well defined measurable physical quantities, the models of GRNs describe dynamics on a spatial scales where the first principle models do not apply. For this reason neither the functional form is fixed, nor the parameters are known. The parameter measurement for such descriptive models is challenging since the parameter is often only defined in the context of the model. Given these challenges it is imperative to devise modeling approaches that deal with parameter uncertainty and describe the dynamics of the GRN in a way that describes the diversity of GRN dynamics across parameters.

In this paper we compare two such complementary approaches. On one hand RACIPE [21] judiciously samples parameters for a reasonable Hill function model of GRN and uses this ensemble as a description of network dynamics. On the other hand, the DSGRN [5] description of the parameter space using all collections of monotone Boolean functions compatible with the network dynamics can be related to switching ODE models and allows precise descriptions of parameter regions with given dynamics by explicit inequalities. Each of these approaches has its limitations. Sampling parameters in high dimensional parameter spaces may miss some dynamics and RACIPE does not come with a guarantee of completeness of represented dynamics in the set of samples. On the other hand, DSGRN can be directly linked only to models with very high Hill coefficient *n* → ∞, which are not biologically realistic. However, DSGRN inequalities do give a detailed functional/phenotypic description of the parameter space, which is lacking in RACIPE. For a given GRN, DSGRN is able to clearly separate the boundaries of the parameter domains, where each domain defines a unique phenotype (mono/bistability, nature of the steady states obtained) emergent from the GRN. Hence, measuring the applicability of such descriptions to RACIPE could lead to an efficient mapping of the phenotypes to the corresponding parameter regimes of biological systems governed by GRNs.

We compared these methods to see if we can combine their advantages and mitigate disadvantages. We established a translation between RACIPE and DSGRN models, allowing us categorize RACIPE sampled parameter sets into DSGRN parameter domains. As expected, we find very good agreement between the phenotypes obtained from DSGRN and RACIPE with high Hill coefficients. Somewhat surprisingly, we also find a very good agreement for low ranges of *n* ∈ [1, 6], specifically in predicting monostability. The bistable parameter nodes from DSGRN always exhibited a mix of mono and bistability in RACIPE. We further found that RACIPE’s sampling is strongly biased towards multistable/oscillatory parameter domains of DSGRN. Despite the differences, we found a striking similarity between the default RACIPE phenotypic distribution and that obtained from DSGRN. This agreement suggests that one can use explicit combinations of parameters supplied by DSGRN to predict emergent phenotypes for ODE models with realistic Hill coefficients.

An important message from this study is the contribution each of the methods can make to enhance the capability of the other. Switching systems, being an approximation of DSGRN, can identify the possibility of multistability for parameter sets with higher accuracy than RACIPE, along with the nature of the states/phenotypes. The approximation of DSGRN output by switching systems is especially useful for networks of higher complexity, where the computation of DSGRN parameter nodes can be computationally expensive.

## 4 Methods

### 4.1 Random Circuit Perturbation (RACIPE)

RACIPE [21] is a tool used to simulate continuous dynamics of gene regulatory networks (GRNs). For a given GRN, RACIPE constructs a set of ODEs representing the interactions in the network. For a node *i*, let *A*_*i*_ and *I*_*i*_ denote the set of all activating and inhibiting input nodes to *i* respectively and *E*_*i*_ denote the expression of node *i*. The dynamics of node *i* is given by the ODE

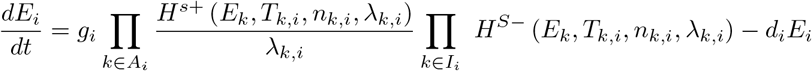

where for node *i, T*_*k,i*_ is the threshold value for the edge from *k* to *i, g*_*i*_ is the production rate, *d*_*i*_ is the degradation rate, *n*_*k,i*_ is the Hill coefficient, *λ*_*k,i*_ is the fold change. The function *H*^*s*+*/−*^ is called the shifted

Hill function, given by

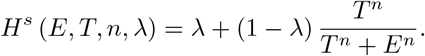

RACIPE simulates this set of ODEs by uniformly sampling parameter sets from a pre-determined range of parameters. These parameter ranges were estimated from BioNumbers [28]. For each parameter set, the ODEs are simulated using the Euler method for multiple initial conditions. The parameters used for each set of simulations and the stable states obtained towards which individual trajectories converge are recorded as an output of the simulation. We then verify the stability of the outputs via the following condition:

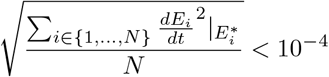

where N is the number of nodes in the network.

### 4.2 Processing RACIPE data

#### 4.2.1 Discretization of RACIPE steady states

For each parameter set sampled by RACIPE, the different initial conditions generated converge to potentially several steady states. Each steady state obtained from RACIPE is assigned a weight, equal to the fraction of initial conditions that converge to the steady state for the corresponding parameter set. This results in a *M* × (*N* + 2) table, where *N* is the number of nodes in the network and *M* is the number of steady state-parameter set combinations. Since each parameter can have more than one steady state, *M >*= *N*. The first column showing the parameter ID, next *N* columns showing the expression level of each node of the network and the last column showing the weight for the corresponding steady state. The node expression values are converted to weighted z-scores by scaling them about their means:

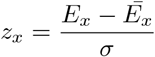

where the steady state expression vector of a node *x* across all steady state-parameter combinations is given by by *E*_*x*_, 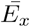 is the weighted mean of the steady state expression of node *x*, and *σ* is the weighted standard deviation of the expression level. The weighted z-scores are then binarised by assigning a value of 1 for positive and 0 for negative weighted z-scores respectively. Hence, each steady state is encoded as a string of zeroes and ones of length that is the the number of nodes in the network.

### 4.3 DSGRN description

A regulatory network summarizes the activating and repressing effects of molecular species on each other. A classical way to model the dynamics associated to a regulatory network is to introduce an ODE system for the concentrations of the various molecular species. A common model is called a *switching system*, in which each regulation event is regarded as a discontinuous and instantaneous switch when a concentration crosses a threshold. These systems are an approximation of the ODE model with graded responses but admit a rather more comprehensive analysis of its solutions [17, 11, 7, 1, 30, 23].

All ODE systems are dependent on a collection of parameters. In [5], we showed that switching systems induce a finite decomposition of parameter space into semi-algebraic regions. Each region represents a *DSGRN parameter* that contains all the information needed to construct a *state transition graph* (STG) that captures a coarse description of the dynamics in phase space of the switching system. This graph is finite, and it is the same for all parameters in the region represented by the DSGRN parameter. This means that the analysis of the dynamics of the switching system over all of the parameter space is computable.

The DSGRN approach collects all of the combinatorial parameters into a graph and computes the STG for each on [5] A condensed version of the STG called a *Morse graph* is a summary of the global structure of the dynamics for each combinatorial parameter. A *Morse node* of the Morse graph is a node representing a strongly connected path component or recurrent component of the STG, and it can be annotated with summary information about the component. We now describe these concepts in greater detail.

A *regulatory network* **RN** = (*V, E*) is a graph with network nodes *V* = {1, 2, …, *n*} and signed, directed edges *E* ⊂ *V* × *V* × {→, ⊣}. For *i, j* ∈ *V*, we will use the notation (*i, j*) ∈ *E* or *i* ⊸ *j* to denote a directed edge from *i* to *j* of either sign; *i* → *j* will denote an *activation* or positive interaction, and *i* ⊣ *j* will denote a *repression* or negative interaction.

We define the *targets* of a node *i* as **T**(*i*) := {*j* | (*i, j*) ∈ *E*} and the *sources* of a node *i* as **S**(*i*) := {*j* | (*j, i*) ∈ *E*}.

A switching system takes the form

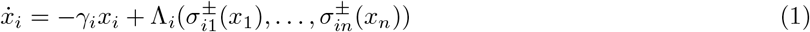

where each argument of Λ_*i*_ is either an increasing or decreasing step function, *σ*^+^ and *σ*^*−*^ respectively, defined by

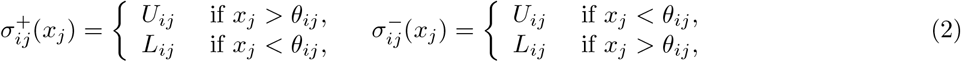

where 0 *< L*_*ij*_ *< U*_*ij*_ represent *lower* and *upper* activation (repression) levels of gene *i* by gene *j* and *θ*_*ij*_ is the *threshold* for the regulatory activity of gene *i* induced by gene *j*. Note that 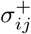 mediates an up-regulation of *I* by *j*, while 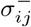 mediates a down-regulation. We assume that all thresholds *θ*_*ij*_ are distinct, which is a generic assumption. The functions Λ_*i*_(*y*_1_, …, *y*_*n*_), *i* = 1, …, *n* are of the form

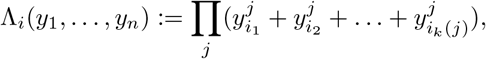

where each *y*_*s*_ occurs exactly once. Such a function is *multi-linear* i.e. linear with respect to each *y*_*i*_, and has all coefficients equal to 1.

The collection of non-negative numbers *γ*_*i*_, *θ*_*ij*_, *L*_*ij*_, *U*_*ij*_ parameterizes the collection of systems (1). Let 𝒫 be the collection of *regular* parameters, i.e. those that satisfy

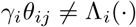

for all (*i, j*) ∈ *E*. Since Λ_*i*_(·) has only finite number of values, 𝒫 is a generic subset of all non-negative parameters.

#### State transition graph

The ordinary differential equation (ODE) system (1) admits a discrete description of phase space that is a directed graph called the *state transition graph*. The assumption that thresholds are distinct implies that each variable *x*_*i*_ has |*T* (*i*)| thresholds. The continuous phase space *R*^*n*^ consists of 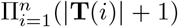 domains *D*(*α*) where *α* is a multi-index with *α*_*i*_ ∈ {0, 1, 2, …, |**T**(*i*)|}. Let

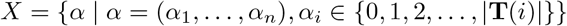

be a generalized hypercube consisting of nodes with labels *α*. Each node represents a corresponding domain *D*(*α*) ⊂ *R*^*n*^.

We construct a state transition graph (STG) on *X* using the fact that solutions in each domain *D*(*α*) converge toward its associated *target point T* (*α*)) = (Λ_1_(*α*)*/γ*_1_, …, (Λ_*n*_(*α*)*/γ*_*n*_). The STG is a graph representation of the *asynchronous update* of the discrete-valued function Λ : *D*(*α*) → *T* (*α*). This means that the STG is a graph on *X* with edges between nodes *α, β* with |*α* − *β*| ≤ 1 assigned in the following way

- if *T* (*α*) ∈ *D*(*α*) then there is a self-edge *α* → *α*;
- if |*α* − *β*| = 1, *α*_*i*_ = *β*_*i*_ − 1 and the threshold between *D*(*α*), *D*(*β*) is *θ*, then *α* → *β* if *T*_*i*_(*α*) *> θ*; and *β* → *α* if *T*_*i*_(*β*) *< θ*.

Note that we can view the STG as a multivalued map ℱ : *X* ⇒ *X*. The DSGRN approach [5, 6, 13, 15, 16, 12] compresses the information about the dynamics of ℱ by computing the strongly connected path components of the STG and the reachability conditions between them. The strongly connected path components are the nodes of a *Morse graph*, which is a Hasse diagram of a partial order imposed by the reachability in the STG. For the purposes of this paper we will be most concerned with Morse nodes that represent a single node in the STG (and thus a single domain in the phase space) where all edges point inwards. This signifies existence of a stable steady state for the ODE (1) which will be denoted as *FP* (*β*) where *β* ∈ *H* determines osition of this steady state with respect to thresholds in the phase space. For instance for the toggle switch, which has one thresholds in each component the space *H* = {0, 1} × {0, 1} and thus the fixed points *FP* can have signatures (00), (01), (10), (11).

#### Parameter space decomposition

The most important construction is the decomposition of the regular parameter space 𝒫 into a finite set of domains, such that for all parameters *p* in one of these domains, the STG is the same.

Let **RN** = (*V, E*) be a regulatory network and let *p* ∈ 𝒫. Then for every *i*, every *j*_*n*_ ∈ **T**(*i*) and every domain *D*(*α*) exactly one of the following inequalities holds

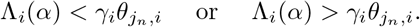

This collection of abstract inequalities, together with the threshold order 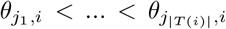 defined by *p* defines a region ℛ of parameter space where every *p* ∈ ℛ induces the same set of inequalities. We call such a collection of inequalities a *DSGRN parameter*.

We can organize domains of parameters in the form of a parameter graph. with a node in a parameter graph (PG), we complete the construction of PG by assigning edges to pairs of nodes that correspond to domains that share a codimension-1 boundary in 𝒫.

Given **RN** = (*V, E*) we represent each DSGRN parameter as a *parameter node* in a DSGRN *parameter graph* **PG**. Two nodes are connected by undirected edge if there is a single inequality change between the corresponding collection of inequalities.

The parameter graph is a product of undirected *factor graphs*, **PG** = Π_*i*_**PG**_*i*_, one associated to each vertex *i* ∈ *V*. The class of possible factor graphs is determined by the topology of network node *i* and the algebraic expression Λ_*i*_ [5].

Given a node 𝒩 ∈ **PG**, each *p* ∈ 𝒩 defines the location of the target points for each of the domains *D*(*α*). Importantly, this location is independent on choice of *p* ∈ N. Therefore the state transition graph and Morse graph can be associated to a parameter node 𝒩 ∈ **PG**.

While for a precise general definition of the parameter graph we refer the reader to [5, 6, 13], we will describe here its construction for the 2-node network examples needed in this paper. For node *i* with one input *i* − 1 and one output *i* + 1, the value of Λ_*i*_ is either *L*_*i,i−*1_ or *U*_*i,i−*1_. Then there are three choices

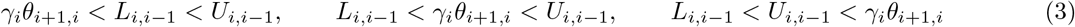

that form the parameter factor graph *PG*_*i*_. Note that in all the cases of Toggle Switch, Dual Activation, and Negative Feedback loop there are two nodes in the network and both nodes have one input and one output. Therefore the parameter graph is a product *PG* = *PG*_1_ × *PG*_2_ with 9 nodes depicted in **Fig 4**. Note that in the horizontal direction, i.e along the *PG*_1_ in the product, the inequalities (3) with *i* = 1 are varying, while in the vertical direction along *PG*_2_ the inequalities (3) with *i* = 2 are changing.

### 4.4 Basin of attraction boundaries for bistable Toggle Switch

In this section we derive analytical expression for stable manifolds of the saddle point in TS based on a switching ODE system.

We consider switching system model of TS with equal decay rates. If we need more general formulation, the below argument can be modified. The model is

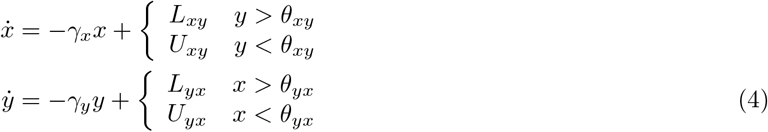

The bistability region in the parameter space satisfies

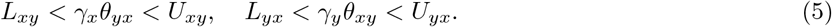

There are four domains in the state space (counterclockwise from bottom right)

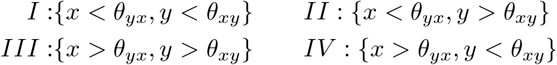

Domain II contains equilibrium FP(0,1) and IV contain equilibrium FP(1,0). We will consider domains I and III which are divided to domains of attraction of two equilibria.

#### Domain III

The equations in this domain are

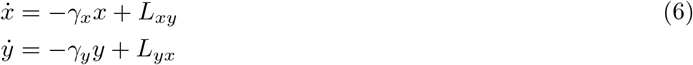

and the solution is

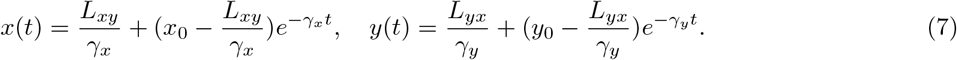

Note that as *t* → ∞ this solution converges to 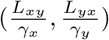 which by the choice (5) of the parameter region, belongs to domain I. Therefore for each initial condition (*x*_0_, *y*_0_) in domain III we can compute time *T*_*xy*_(*x*_0_, *y*_0_) where this solution crosses *θ*_*xy*_ and the time *T*_*yx*_(*x*_0_, *y*_0_) where it crosses *θ*_*yx*_. Importantly, if *T*_*xy*_(*x*_0_, *y*_0_) *< T*_*yx*_(*x*_0_, *y*_0_) then (*x*_0_, *y*_0_) is in the domain of attraction of FP(1,0) in domain IV, and if *T*_*xy*_(*x*_0_, *y*_0_) *> T*_*yx*_(*x*_0_, *y*_0_) then (*x*_0_, *y*_0_) is in the domain of attraction of FP(0,1) in domain II. Consequently, the condition *T*_*xy*_(*x*_0_, *y*_0_) = *T*_*yx*_(*x*_0_, *y*_0_) defines the boundary between these domains of attraction.

The equations for *T*_*xy*_(*x*_0_, *y*_0_) and *T*_*yx*_(*x*_0_, *y*_0_) are

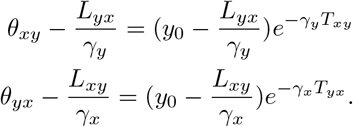

This gives

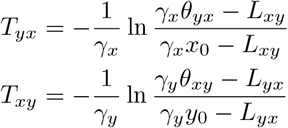

This leads to the following equation for the separatrix between two domains of attraction in domain III:

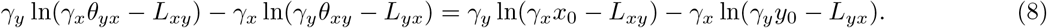

Notice that the left hand side is a constant, while the right hand side depends on the values (*x*_0_, *y*_0_). Therefore domain of attraction within domain III of the fixed point in domain IV (**FP(1**,**0)**) will satisfy

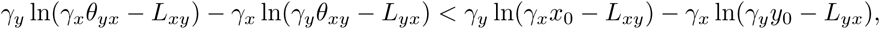

while the domain of of the fixed point in domain I I will have the inequality reversed.

#### Domain I

The equations in this domain are

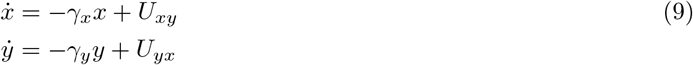

and the solution is

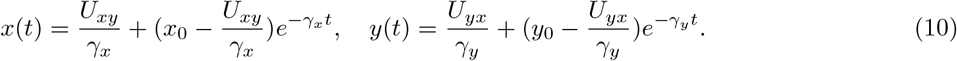

These equations are analogous to (10) with constants *L*_*xy*_, *L*_*yx*_ replaced by *U*_*xy*_, *U*_*yx*_, respectively. Therefore, in analogy with (8), we have the following equation for the separatrix between two domains of attraction in domain I:

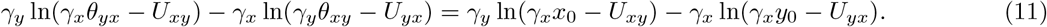

We use equations (8) and (11) to classify RACIPE-selected initial conditions (*x*_0_, *y*_0_) as belonging to the basin of attraction of either of the stable equilibria in domains II and IV.

Similar equations can be written for switching system of DA, where domain I contains FP(0,0) and domain III contains FP(1,1). The corresponding equations defining the boundaries of the basins of attraction are:

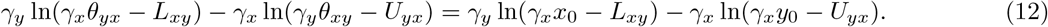

for domain I and

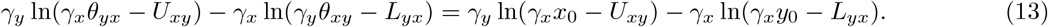

for domain III respectively.

### 4.5 Translating DSGRN to Shifted Hill functions

The RACIPE model for the dynamics of a node *i* is given by

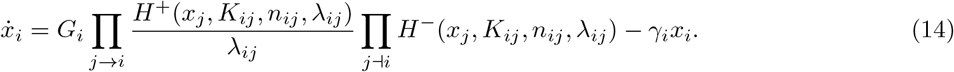

Here, *H*^*±*^ are shifted Hill funtions, given by

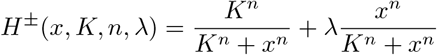

with the requirement that *λ >* 1 for *H*^+^ and *λ <* 1 for *H*^*−*^. Shifted Hill functions are monotone sigmoid functions. In particular, as *n* → ∞, they converge to switching functions. We note that *H*^*±*^(0) = 1 and

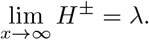

It follows that the range of *H*^+^ is (1, *λ*), and the range of *H*^*−*^ is (*λ*, 1). Since the effect of an activating edge in (14) is 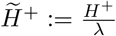, this range of 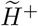 is 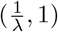. The final parameter of RACIPE formalism is the basal production rate *G*_*i*_ at each network node.

The DSGRN model for the dynamics of node *i*, assuming product interactions between edges with common target node, is given by

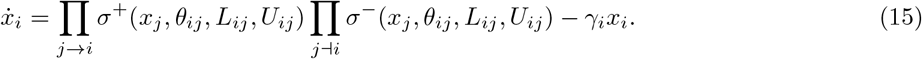

The switching functions *σ*^+^ and *σ*^*−*^ are defined in (2). We first observe that in the limit *n* → ∞ a half-saturation parameter *K*_*ij*_ in *H*^*±*^ of (14) becomes the threshold *θ*_*ij*_ in functions *σ*^*±*^. We therefore equate these parameters in the two models

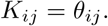

We now address conversion between the fold multiplication parameters *λ*_*ij*_ and basal rate *G*_*i*_ in (14) and parameters *L*_*ij*_ and *U*_*ij*_ in (15). At first glance, there are more parameters in the DSGRN model than in the RACIPE formulations. When the inputs to a node *i* combine as a product, which is the case in both (14) and (15), the key observation is that the highest value of this product at node *i* is *G*_*i*_ in (14) and ∏_*j→i*_ *U*_*ij*_ in (15).

Therefore a natural correspondence is

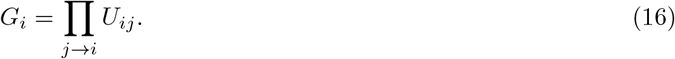

#### Converting a DSGRN Model to a RACIPE Model

With the choice above the remaining conversions are as follows. Given DSGRN model, we first compute the basal rate *G*_*i*_ for each *i* from (16). Let *m*_*i*_ be the number of incoming edges to node *i*. Then the conversion between *L*_*ij*_ and *λ*_*ij*_ is as follows:

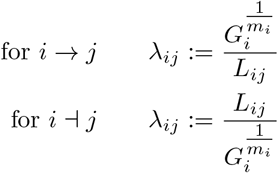

In addition, since DSGRN corresponds to a limit as Hill coefficients approach infinity, the Hill coefficient of RACIPE is *n*_*ij*_ = ∞ for each *i, j*.

#### Converting a RACIPE Model to a DSGRN Model

Given a RACIPE model, we choose to evenly attribute the basal production rate, *G*_*i*_, to each input edge by setting

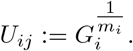

Let *m*_*i*_ be the number of inputs to node *i*. The corresponding DSGRN parameters are given by

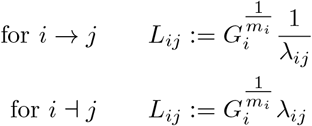

With this choice of correspondence in parameters each shifted Hill function in the RACIPE model converges to the corresponding switching function in the DSGRN model as *n* → ∞. A consequence of Theorem 3.11 of [10] is that the fixed points of the DSGRN model are in one-to-one correspondence to the fixed points of the RACIPE model for large enough *n*.

## Acknowledgement

TG, BC and WD were partially supported by NSF grant DMS-1839299 and NIH 5R01GM126555-01. KH was supported by the Prime Minister’s Research Fellowhip. MKJ was supported by Ramanujan Grant SB/S2/RJN-049/2018. We acknowledge the Indigenous nations and peoples who are the traditional owners and caretakers of the land on which this work was undertaken at Montana State University.

---

**Figure S1:**
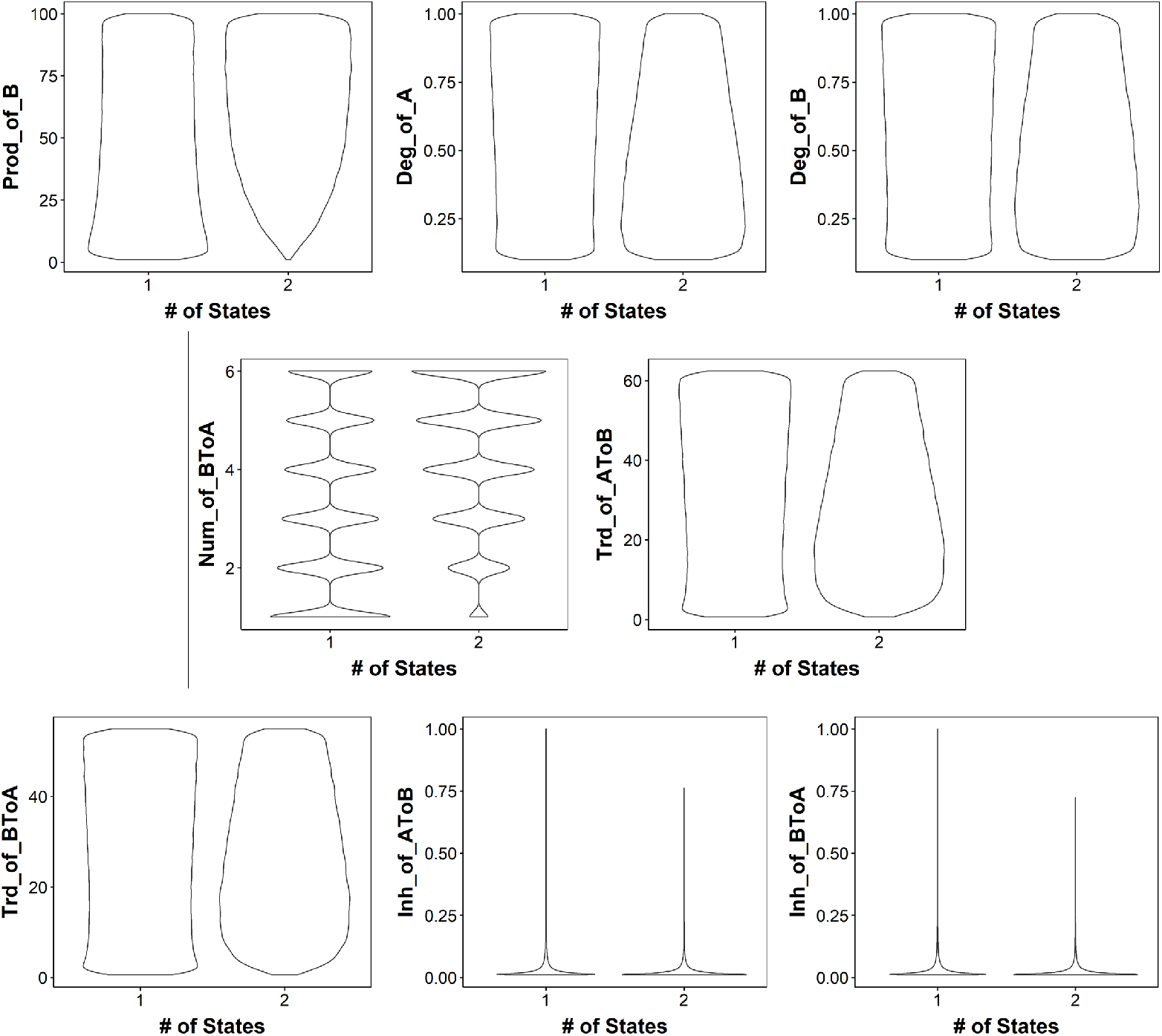
Parameter distribution for monostable and bistable parameter sets for TS in RACIPE

**Figure S2:**
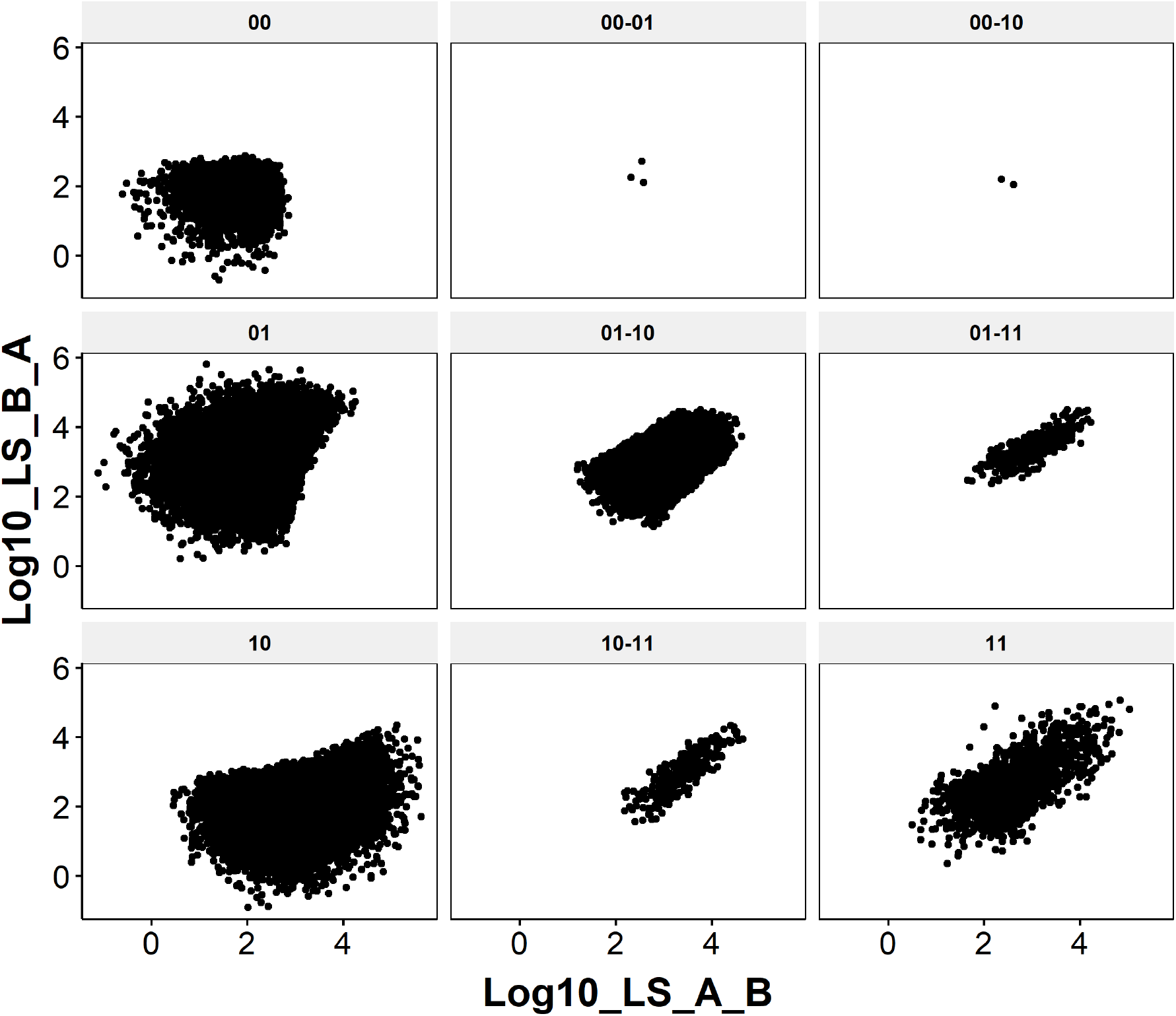
Scatterplot showing the Link strength values for parameters corresponding to different phases.

**Figure S3:**
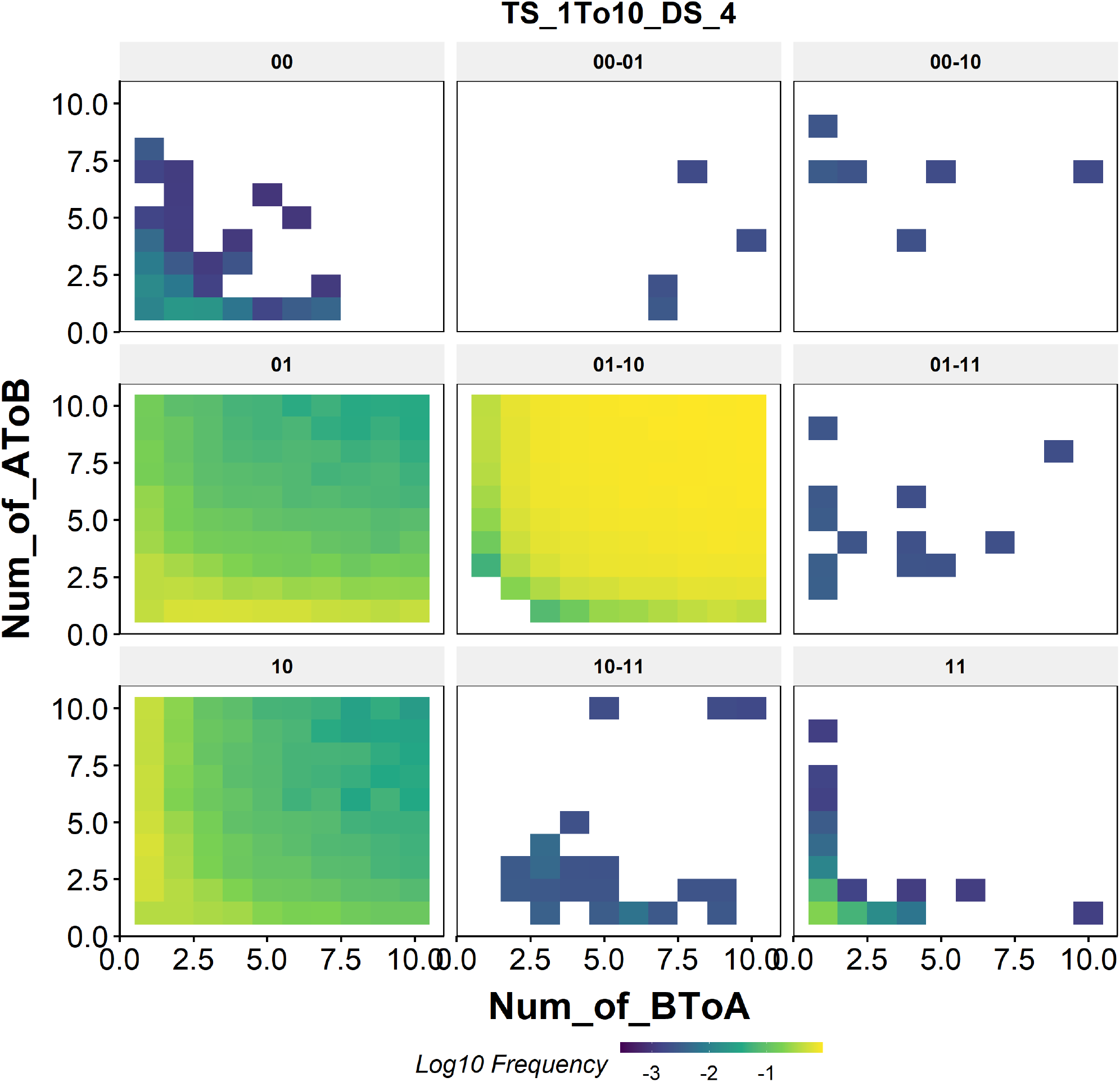
Dependence of phase frequency of TS in parameter node 4 (bistable parameter node) on Hill coefficients in the 1-10 range. The color denotes, in log scale, the frequency of a phase for a given combination of Hill coefficients.

**Figure S4:**
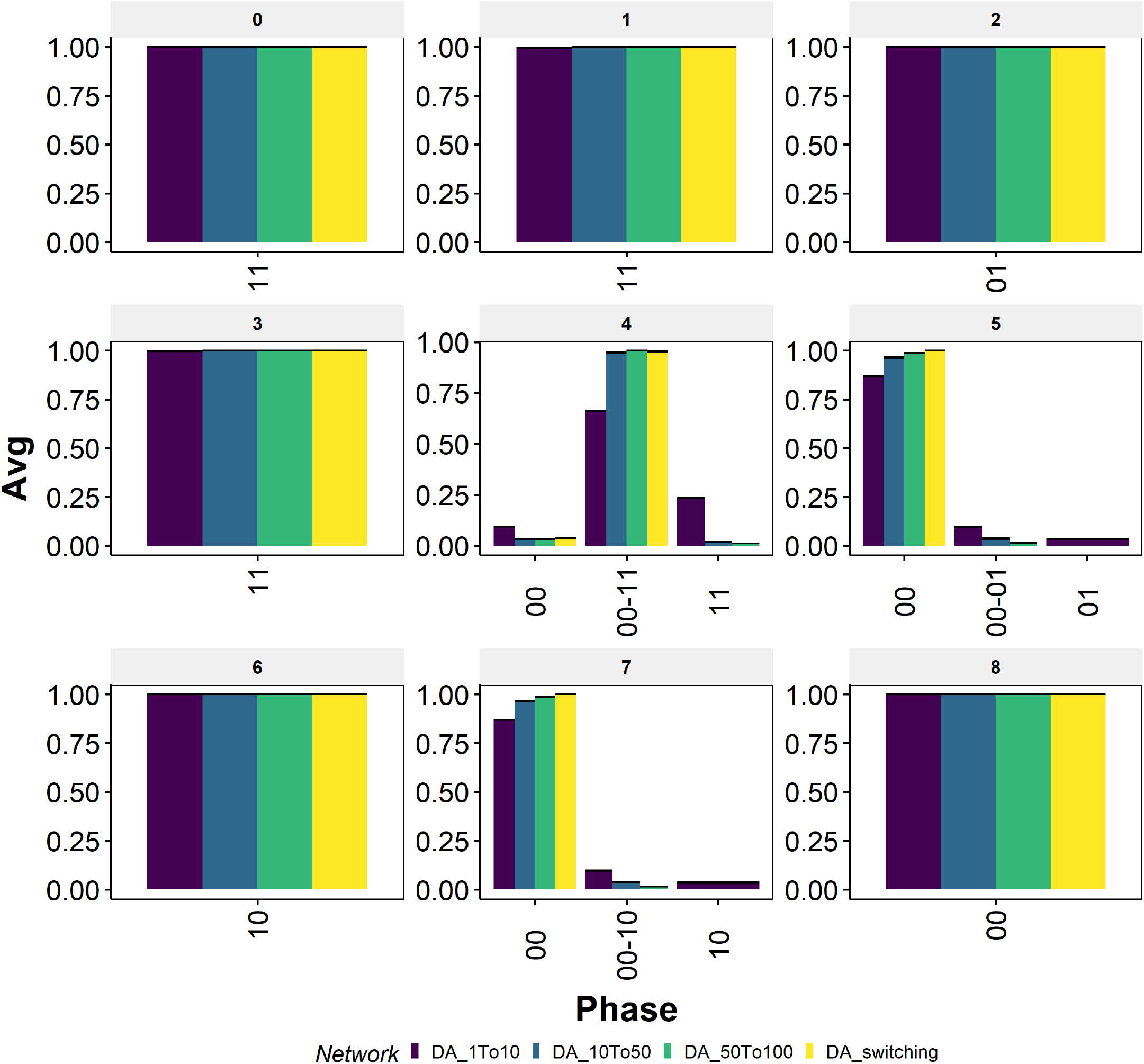
Phase distribution of switching system against different ranges of Hill coefficients in RACIPE for DA. The switching system is represented by yellow colored bars. Default RACIPE conditions are represented by the dark blue colored bars. The range of Hill coefficients in each case is reported in the color-legend.

**Figure S5:**
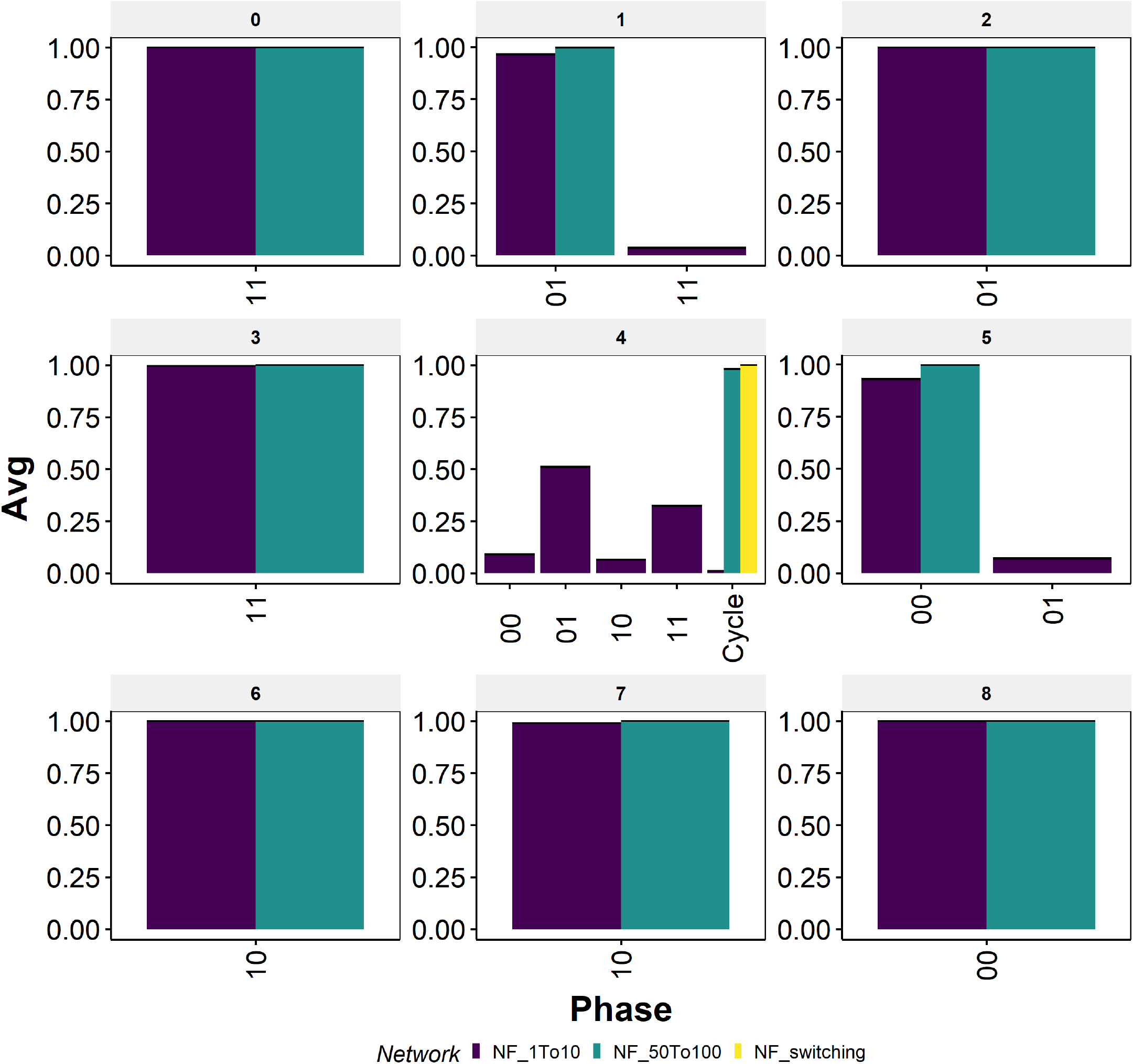
Phase distribution of switching system against different ranges of Hill coefficients in RACIPE for NF. The switching system is represented by yellow colored bars. Default RACIPE conditions are represented by the dark blue colored bars. The range of Hill coefficients in each case is reported in the color-legend.

**Figure S6:**
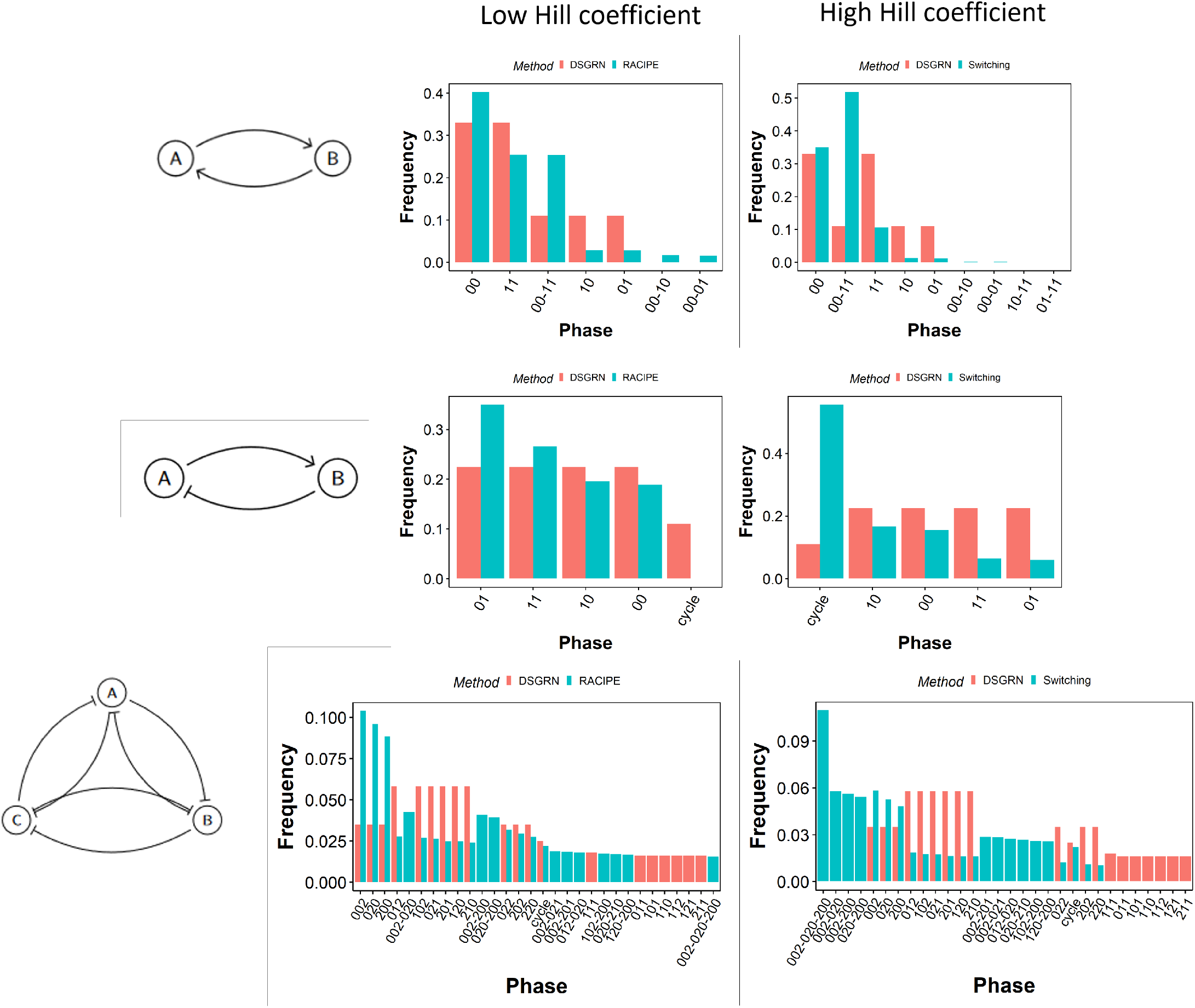
Comparison of the DSGRN phase distribution with that of RACIPE at low and high hill coefficients for DA, NF and TT.

